# Weevil borers affect the spatio-temporal dynamics of banana Fusarium wilt

**DOI:** 10.1101/2021.04.01.437979

**Authors:** Daniel Heck, Gabriel Alves, Eduardo S. G. Mizubuti

## Abstract

Dispersal of propagules of a pathogen has remarkable effects on the development of epidemics. Previous studies suggested that insect pests play a role in the development of Fusarium wilt (FW) epidemics in banana fields. We provided complementary evidence for the involvement of two insect pests of banana, the weevil borer (*Cosmopolites sordidus* L. - WB) and the false weevil borer (*Metamasius hemipterus* L. - FWB), in the dispersal of *Fusarium oxysporum* f. sp. *cubense* (*Foc*) using a comparative epidemiology approach under field conditions. Two banana plots located in a field with historical records of FW epidemics were used, one was managed with *Beauveria bassiana* to reduce the population of weevils, and the other was left without *B. bassiana* applications. The number of WB and FWB was monitored biweekly and the FW incidence was quantified bimonthly during two years. The population of WB and the incidence (6.7%) of FW in the plot managed with *B. bassiana* were lower than in the plot left unmanaged (13%). The monomolecular model best fitted the FW disease progress data and, as expected, the average estimated disease progress rate was lower in the plot managed with the entomopathogenic fungus (*r* = 0.0024) compared to the unmanaged plot (*r* = 0.0056). Aggregation of FW was higher in the field with WB management. WB affected the spatial and temporal dynamics of FW epidemics under field conditions and brought evidence that managing the insects may reduce FW of bananas intensity.

## INTRODUCTION

Understanding the ways a pathogen is dispersed is one of the most important tasks in the epidemiology of plant diseases. For Fusarium wilt (FW) of bananas, caused by *Fusarium oxysporum* f. sp. *cubense* (*Foc*) (E. F. Smith) Snyder and Hansen, there is limited information about the mechanism of pathogen dispersal available. It is suggested that *Foc* can be dispersed mainly by human influence, such as exchange of asymptomatic propagation material, cultural practices performed with infested tools or conducted by untrained workers, and movement of soil particles adhered to boots, machinery and tires of vehicles (Dita et al. 2018; Ploetz et al. 2015). Good management practices, sterilization of materials, and appropriate training of plantation workers can greatly reduce dispersal and the likelihood of the pathogen introduction in areas without records of FW.

Even under conditions of proper management practices, severe epidemics can occur. Natural dispersal agents or processes such as wind, water, animals, including mammals and insects, are more difficult to detect and control. Some reviews highlighted the potential of these dispersal processes in the movement of *Foc* propagules at short or long distances (Dita et al. 2018; Ploetz et al. 2015; Ploetz 2015; Ghag et al. 2015). Surface water and rivers infested with *Foc* propagules were suggested as a cause of the rapid expansion of FW epidemics in China, Malaysia, and the Philippines (Ploetz et al. 2015; Xu et al. 2003). Fungal species, as *F. oxysporum*, do not move long distances in soil without host tissues. It is expected that the main way of pathogen dispersal in the environment occurs when the pathogen colonizes roots of susceptible hosts or is passively transported by animals, wind, and water. Inoculation normally occurs when root growth contacts the inoculum distributed in the soil or by root-to-root contact (Rekah et al. 1999). Eventually, *Foc* may be dispersed aerially. The presence of sporodochia and hyphal growth externally in plant tissues suggests aerial dispersal of *Foc* (Warman and Aitken 2018).

Any animal or material that can carry soil particles in banana fields is a potential dispersal agent of *Foc*. The free circulation of animals such as feral pigs in banana fields was cited as a potential mechanism of dispersal of *Foc* (Biosecurity of Queensland 2016). The potential role played by the banana weevil borer (*Cosmopolites sordidus* L., Coleoptera: Curculionidae), referred here as weevil borer (WB), in special, was suggested to be responsible for the inoculation of isolated banana plants in the field (Meldrum et al. 2013). Viable inoculum of *Foc* was recently reported in the external and internal body of WB, and the inoculum remained viable up to 3 days (Sánchez et al., 2021). Weevil borer is the main insect pest in banana fields and can occur in large populations (Gold et al. 2001). Other pests, such as the West Indian sugarcane weevil (*Metamasius hemipterus* L., Coleoptera: Curculionidae), referred to as false weevil borer (FWB) by banana growers, is an important pest in sugarcane and many crops. The FWB is also observed causing damages in plantain fields under high populations (Fancelli et al. 2012). Curculionidae members acting as vectors of plant diseases are commonly reported in many crops. *M. hemipterus* and *Rhynchophorus palmarum* (Coleoptera: Curculionidae) were suspected to be vectors of the nematode *Bursaphelenchus cocophilus* (*=Rhadinaphelenchus cocophilus*), the causal agent of red ring disease in oil palm (Mora et al. 1994; Hagley 1963). *R. palmarum* was also reported as dispersal agent of bud rot disease (*Phytophthora palmivora*) in oil palm (Plata-Rueda et al. 2016) and stem bleeding (*Thielaviopsis paradoxa*) in coconut palm (Carvalho et al. 2011).

Since the study carried out by Meldrum et al. (2013) the effective contribution of banana weevil in the FW epidemics under field conditions remains unanswered. Nevertheless, these answers may be useful to the development of preventive measures against quarantine pathogens, as *Foc* Tropical Race 4 (TR4), and effective management strategies to control the expansion of FW foci within the field. The objective of this study was to understand the relationship of two potential vectors of *Foc*, *C. sordidus* and *M. hemipterus*, in the FW epidemics using a comparative epidemiology approach: a spatio-temporal study under field conditions.

## MATERIAL AND METHODS

### Field

One banana field where FW was known to have occurred was selected to study the association between insect pests and FW epidemics. The field was located in the municipality of Teixeiras, Minas Gerais state, Brazil, cultivated with ‘Prata’ type banana (Pome subgroup, AAB) and managed as a low-input system. The field was subdivided into two plots. In one plot (20° 38’ 21.444’’ S; 42° 50’ 1.752’’ W and 772 masl) of 1.03 ha, low population of WB was attempted by means of one initial application of a chemical insecticide followed by several applications of a biological control agent. In the second plot (20° 38’ 19.5’’ S; 42° 50’ 2.544’’ W and 759 masl) of 1.2 ha no control practices for WB and FWB were adopted. The management of insects was conducted at every three months. Carbofuran (Furadan 50 G^®^, FMC Corporation) was applied manually with 3 g of commercial product per trap and 40 traps per ha at the beginning of the experiment. Subsequent management practices to control insects were performed with *Beauveria bassiana* (Ballvéria WP^®^, Ballagro; and Beauveria Oligos WP^®^, Oligos Biotec) applied every three months at a rate of 5 g per trap and 30 to 40 traps per ha. Traps were made by cross-cutting pseudostems of harvested plants at approximately 40 cm from the soil. The biological insecticide was applied on the flat cut surface of one half of the trap, and then the above section was placed back on top of the treated half. In addition to the chemical and biological control methods, insects collected in the monitoring traps were eliminated every two weeks. The chemical, biological and manual control of the insects were only performed on the managed plot.

### Insects monitoring

The second type of pseudostems trap was used to monitor the WB and FWB populations. Pseudostems of harvested plants were cross-cut in sections of 30 cm length and subsequently lengthwise in half such as to produce two hemicylindrical traps. The sections were placed with the cut side down (in contact with soil). Fifteen pseudostem traps per plot were made and placed approximately 15 cm apart from the banana mat. The traps were randomly distributed in the plots and maintained in the same place throughout the experiment. Monitoring was performed from May 2017 to February 2019. Traps were evaluated by counting the number of WB and FWB every two weeks and replaced by new ones. Data were analyzed by the paired t-test on each assessment date with the STATS package (R Core Team, 2018). T-test was also used to analyze differences in the average number of WB and FWB between the plots.

### Fusarium wilt assessment

Banana plants were visually assessed for external and internal symptoms of FW. If external symptoms such as wilt, yellowing, and collapse of older leaves, and/or splitting at the base of pseudostem in at least one plant of the mat was observed, a small cut was made in the pseudostem to inspect for internal symptoms. Internally, diseased plants presented yellow, reddish-brown, or black discolorations of vascular tissues. If external and internal symptoms were present, the mat was considered diseased and was georeferenced using a handheld GPS device (GPSMAP^®^ 64, Garmin). The diseased banana mat was identified with a striped plastic tape to avoid double quantification in future assessments. GPS files with the geographic location of diseased plants and polygons were extracted and converted to text files and imported in R Statistical Software version 3.5.1 (R Core Team 2018). Fragments (*N* = 10; 5 cm of length x 5 cm of width) of pseudostems taken from symptomatic plants from the field were used to isolate the pathogen and confirm the presence of *Foc* by morphological (Leslie and Summerell 2006) and molecular methods (O’Donnell et al. 1998; Heck et al. *in preparation*).

### Spatio-temporal analyses

#### Temporal analysis

The area size, centroid, and total number of plants in each plot were estimated using the polygon data set and the plant spacing information (distance within and between rows). The diseased plant data set was used to estimate the total number and location of diseased plants (*x*) in plots at each assessment. The incidence (*p*) was calculated as *p* = Σ*x*/ Σ*n*, where *x* was the number of diseased plants and *n* the estimated number of total plants (healthy and diseased) on each plot. The incidence was calculated at each assessment date. The Monomolecular, Logistic and Gompertz models were fit to the disease incidence data plotted over time by nonlinear regressions using the nlsLM function from MINPACK.LM package (Elzhov et al. 2016). The choice of the best model was performed by the linear mixed-effects model (Laird and Ware 1982) using the LME function from the NLME package (Pinheiro et al. 2018).

#### Spatial analysis

Two types of spatial analyses were conducted: a point-pattern and a geostatistical-based approach. Field maps with the geolocation of diseased plants were *quadratized* using the QUADRATCOUNT function of the SPATSTAT package (Baddeley and Turner 2005). The quadrat size used was 2×2, i.e. the area occupied by two plants within and between rows, respectively. The number of diseased plants in each quadrat was determined and used to perform the spatial analysis. The exact number of plants inside each quadrat could not be computed because the distance within and between rows was irregular. The estimated precision of the GPS device varied between 3 and 7 m during the assessments depending on the environmental conditions.

Dispersion index for binomial data, *D*, referred sometimes as the ratio between variances was used for the point-pattern approach. A χ^2^ test was performed to test the null hypothesis of dispersion index (*D*) = 1. The analysis was conducted using the AGG_INDEX function from the EPIPHY package (Gigot 2018). Spatial Analysis by Distance IndicEs (*SADIE*) was the method chosen in the geostatistical-based approach. *SADIE* uses the location of the sampling units (i.e. quadrats) and the number of individuals (i.e. diseased plants) inside the unit to analyze the spatial arrangement by the distance to regularity (*D_r_*) criterion. The *D_r_* is achieved when all the sampling units have the same number of diseased individuals. *SADIE* returns an overall aggregation index (*I_a_*) which reflects the ratio between the distance moved to achieve a regular pattern for the observed data and a theoretical mean to regularity based on random permutations. The index developed by Li et al. (2012) was computed by the SADIE function from the EPIPHY package. A random pattern was inferred when the *D* and *SADIE* aggregation indices were equal to 1, an aggregated pattern when they were higher than 1, and regular when less than 1.

#### Spatio-temporal study

The relationship between the observed and theoretical variances of FW incidence per sampling unit over time was evaluated. The equation for a binary power law, *log_10_*(*s^2^*) = *log_10_*(*A*) + *b.log_10_*(*s^2^*_bin_), where *s^2^* is the observed variance, *s^2^_bin_* the theoretical variance assuming a binomial distribution, and *A* and *b* are the parameters to be estimated. Estimates of *A* = *b* =1 suggest a random distribution; if *A* >1 and *b* =1 suggest an aggregated pattern with fixed value independently of the incidence level; and if both parameters, *A* = *b* > 1, the level of aggregation varies with incidence (Madden and Hughes 1995). A t-test was performed to test the null hypothesis for individual plots, and to test if the parameters differed between the plots.

## RESULTS

Management and monitoring of WB were successfully carried out from May 2017 to February 2019. The overall number of FWB collected (median of 2.3 per trap) was significantly higher than WB (median of 1.1 per trap) in each assessment (*P* < 0.001). The number of WB was lower in the field managed with *B. bassiana* (median of 0.67 per trap) when compared to the unmanaged field (1.6 per trap) (*P* < 0.001; Fig 1a). Management with *B. bassiana* did not affect the population of FWB in the same period (*P* = 0.789; Fig 1b).

**Fig 1.**
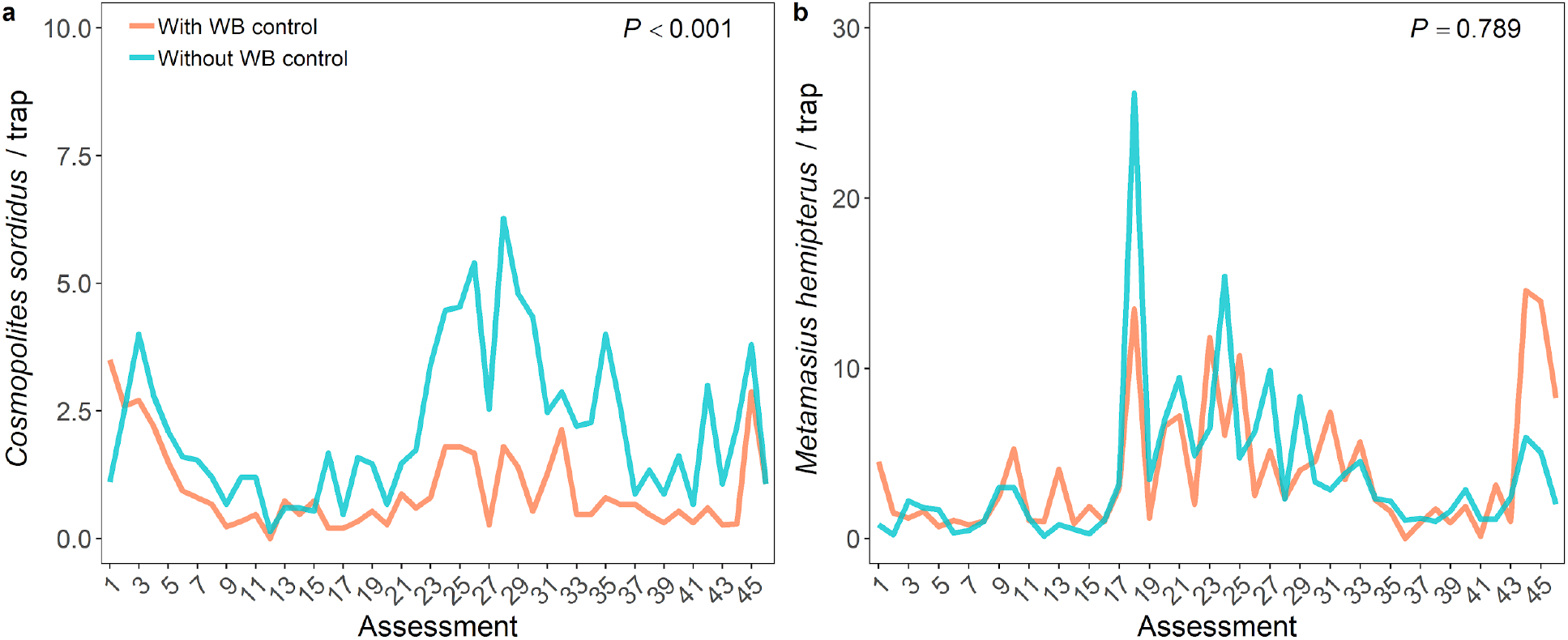
Biweekly monitoring of weevil borer (*Cosmopolites sordidus*, WB) (a); and false weevil borer (*Metamasius hemipterus*, FWB) (b) in fields with and without control of WB, from May 2017 to February 2018.

Fusarium wilt incidence was lowest in the first assessment: 0.9% and 2% in the plots with and without WB management, respectively (Fig 2). These two plots already had diseased plants when the assessments began. After 12 assessments disease incidence in these two plots reached up to 6.7% and 13% in fields with and without WB management, respectively (Fig 2). The monomolecular model described the disease progress in both plots (Table 1). Significant differences were not observed for the initial incidence parameter (*P* = 0.135) between the plots and the disease progress rate was significantly higher in the unmanaged plot (*P* < 0.001; Fig 2).

**Fig 2.**
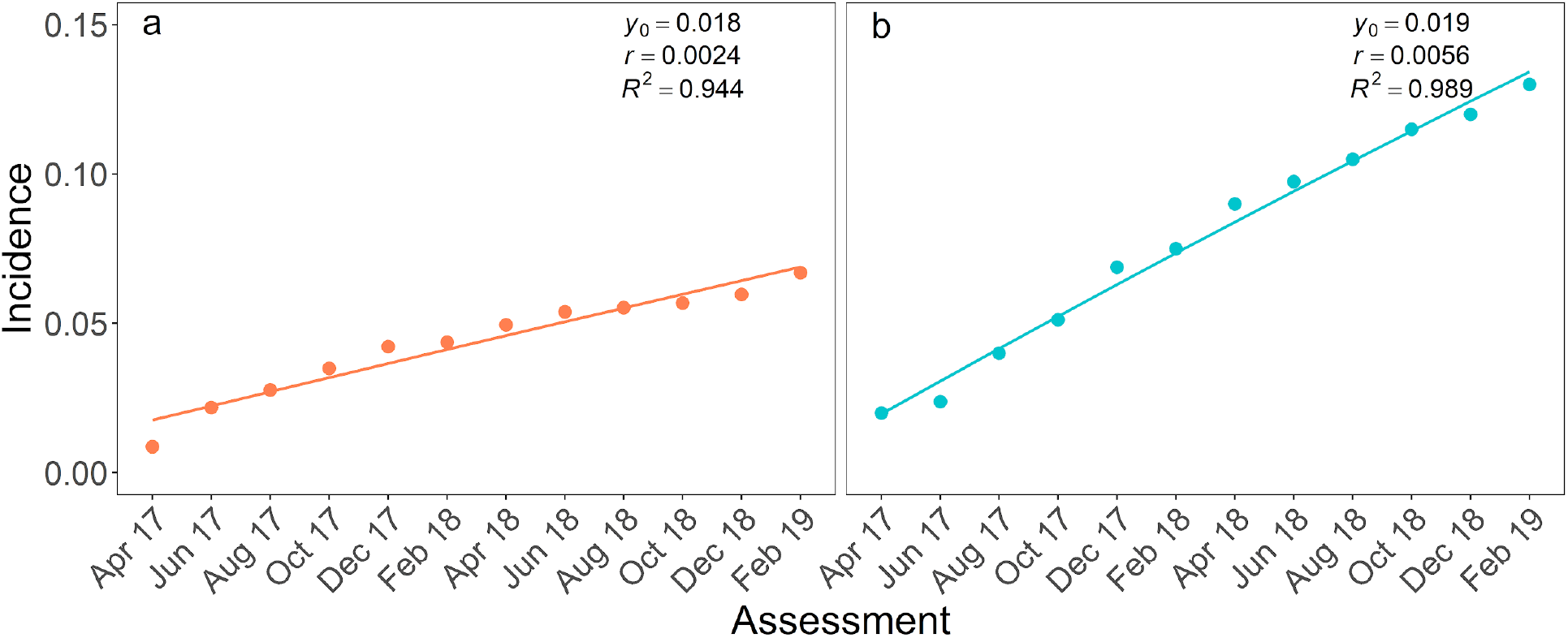
Incidence of Fusarium wilt of bananas in plots with (a) and without (b) management of weevil borer (*Cosmopolites sordidus*, WB) from April 2017 to February 2019 (points). Monomolecular model adjusted to incidence data (curves) The plots were located in the municipality of Teixeiras, Minas Gerais state, Brazil. Estimated initial incidence (*y_0_*), progress rate (*r*) of epidemics, and the measure of the quality of the adjusted monomolecular model (*R^2^*) are presented.

**Table 1.** Summary of statistics used to evaluate the disease progress curve of Fusarium wilt of banana in two plots, with and without weevil borer management, located in municipality of Teixeiras, Minas Gerais state, Brazil, from April 2017 to February 2019.

| Model | AIC | BIC | logLikelihood | Best adjust / *n* |
| --- | --- | --- | --- | --- |
| Monomolecular | -219.95 | -213.98 | 115.98 | 2/2 |
| Logistic | -137.39 | -131.41 | 74.69 | 0/2 |
| Gompertz | -153.44 | -147.47 | 82.72 | 0/2 |

AIC: Akaike Information Criterion; BIC: Bayesian Information Criterion;

Spatial analyses were computed for both fields using point-pattern and geostatistical-based approaches. Aggregation of FW epidemics was inferred in all assessments and both plots by the dispersion index (Fig 3a). Aggregation was lowest in the first two assessments and highest in the last. Aggregation was also higher in the plot with WB management than in the unmanaged field (*P* < 0.001). The dispersion index ranged from 1.90 to 2.47 with a mean of 2.23 in the managed plot and from 1.34 to 2.27 with a mean of 1.86 in the unmanaged (Fig 3a).

**Fig 3.**
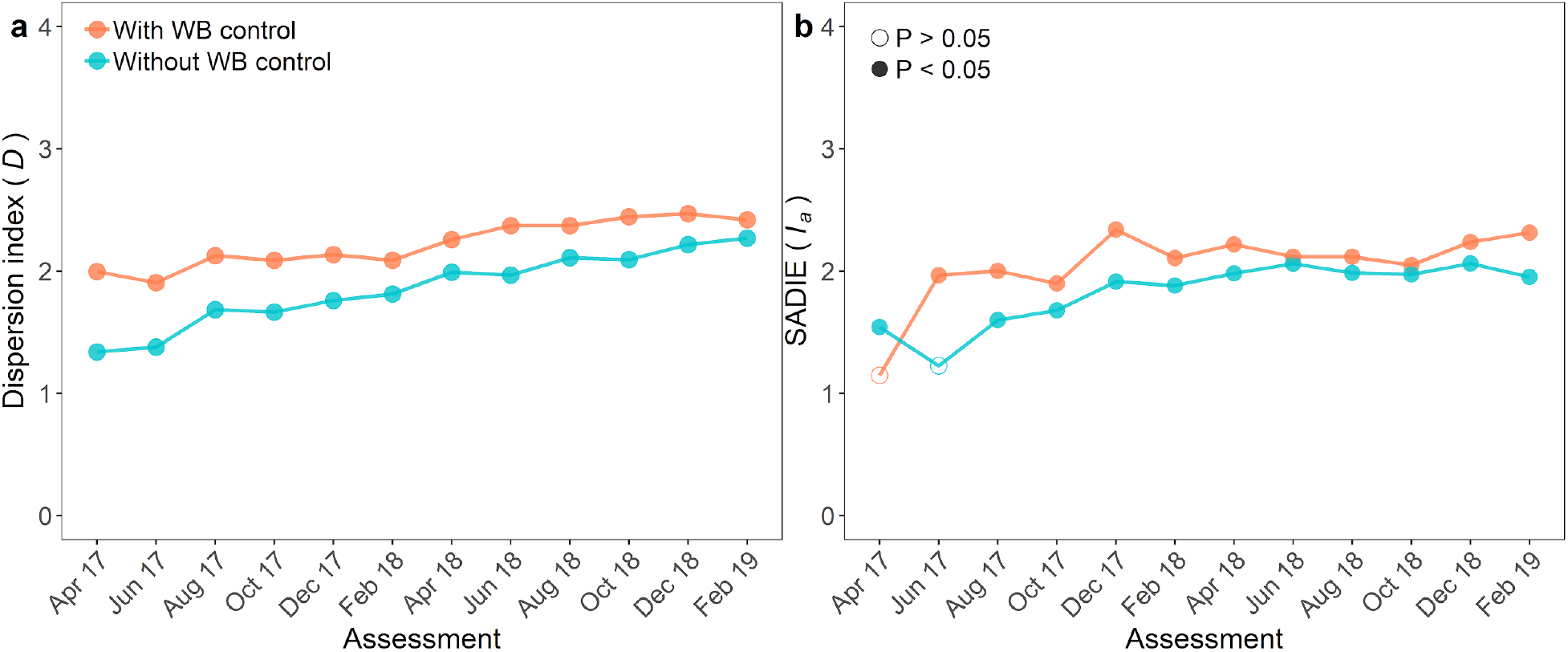
Dispersion index (*D*; a) and aggregation index (*I_a_*) of Spatial Analysis by Distance Indices (*SADIE*; b) of incidence of Fusarium wilt of banana in two plots, with and without management of weevil borer (*Cosmopolites sordidus* - WB), from April 2017 to February 2019. The plots were located in the municipality of Teixeiras, Minas Gerais state, Brazil. *P*-value was represented by empty (*P* > 0.05) or filled (*P* < 0.05) symbols.

The random pattern was detected using *SADIE* in the first two assessments only. In these assessments the *I_a_* values were low (1.15 and 1.23) (Fig 3b). The highest *I_a_* value (2.34) was observed in the managed plot. The degree of aggregation obtained by *SADIE* differed between the plots. The *I_a_* was higher in the managed (median of 2.04) compared to the unmanaged plot (1.82) (*P* = 0.016; Fig 3b).

Binary power law expresses the relationship between the logarithm of the observed variance and the logarithm of theoretical variance assuming a binomial distribution and was well adjusted in plots with (*R^2^* = 0.992) and without (*R^2^* = 0.998) WB management (Fig 4). Parameters of the binary power law, *A* and *b*, were higher than 1 (*P* < 0.001) for both plots. The level of aggregation in plots varied according to the incidence observed. *Log_10_*(*A*) and *b* also differed between plots (*P* < 0.001). The aggregation in the unmanaged plot was more affected by the changes in incidence than in the managed plot (Fig 4).

**Fig 4.**
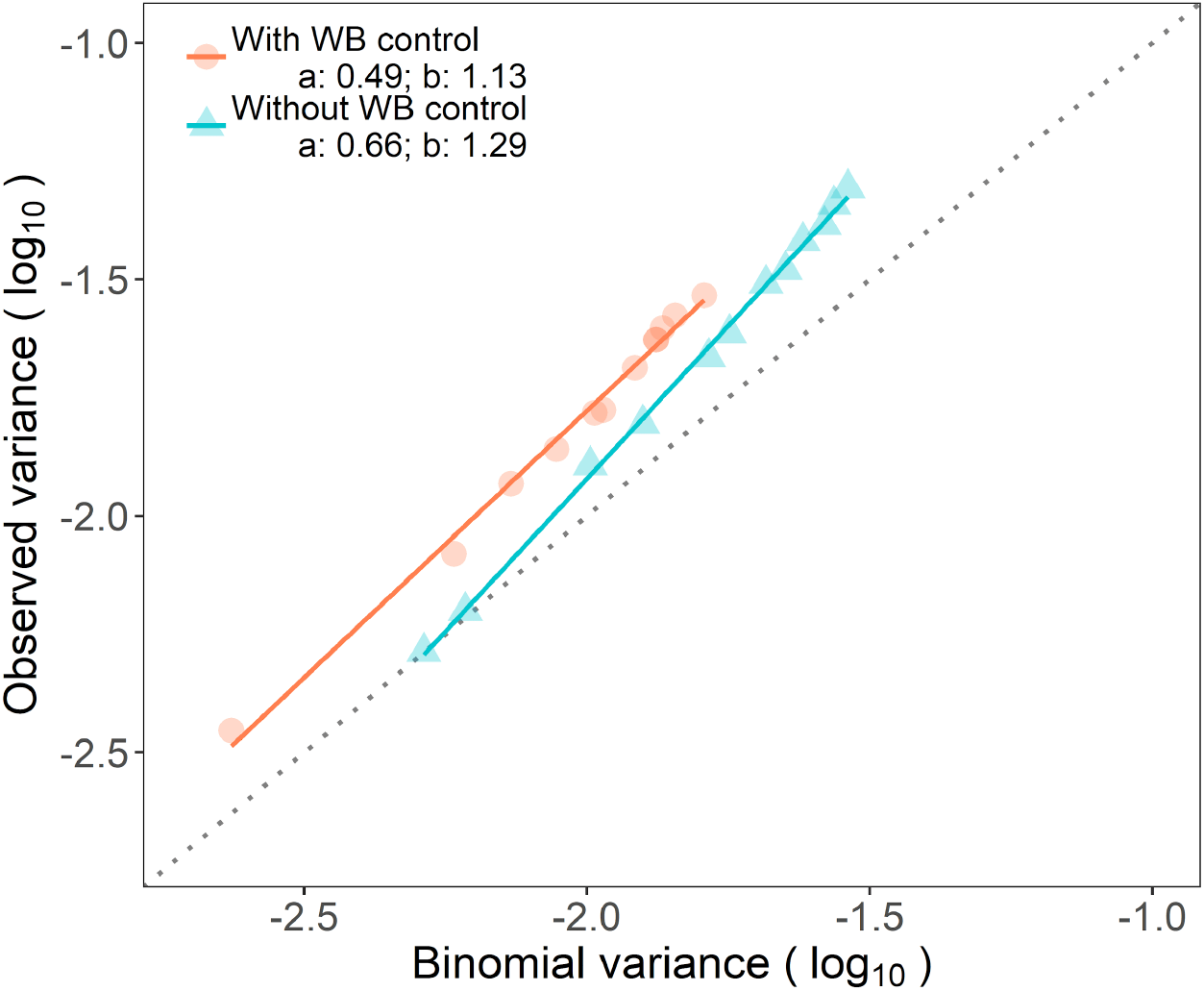
Binary power law of successive assessments for incidence of Fusarium wilt of banana from April 2017 to February 2019 in two plots, with and without management of weevil borer (*Cosmopolites sordidus* - WB). The plots were located in Teixeiras, Minas Gerais state, Brazil.

## DISCUSSION

Currently, there is high interest in the dispersal mechanisms of *Foc* (Dita et al. 2018; Ploetz et al. 2015). This information is crucial for the management of FW in already infested areas as well as for setting exclusion and mitigation actions in disease-free fields. The role played by pests that are present in banana fields still needs to be elucidated. Epidemiological information can be useful to develop management strategies to effectively reduce the spread of *Foc* and reduce the damage caused by FW. In this study, the spatio-temporal approach was used to try to shed light on the putative role played by insect vectors on epidemics of FW.

The management of banana weevil was performed using *B. bassiana*, an entomopathogenic fungus widely used to control banana weevils and false weevils (Kaaya et al. 1993; Akello et al. 2008; Pauli et al. 2011; Mesquita et al. 1981). The entomopathogen enters the insect through the spiracles, digestive system, or insect cuticle and grows in hemocoel and muscle tissues, destroying tracheal taenidia and fat bodies (Kaaya et al. 1993). In both plots, the population of FWB was higher than that of WB. False weevil borer is considered a secondary pest in banana and other crops (Fancelli et al., 2012). However, in high populations, it damages banana plantations and management needs to be considered (Fancelli et al. 2012). In this study, the population of *M. hemipterus* was not affected by the management of the insects after the application of *B. bassiana*, even though many dead individuals were constantly found in traps. FWB has the ability to fly, giving it higher mobility compared to the WB in the fields. In addition, an abandoned banana plantation close to the managed plot probably masked the counts of FWB. These facts could have increased the chances of FWB being attracted to the pseudostems traps (Dolinski and Lacey 2007).

Differences in the disease progress curve of FW incidence were observed between the two plots. The incidence was lowest in the first assessment and increased over time, reaching 13% in the unmanaged and 6.7% in the managed plot. These values can be considered low when comparing with reports in Tanzania and Indonesia where the FW incidence reached 77% and 100%, respectively (Karangwa et al. 2016; Hermanto et al. 2011). However, the cultivars, environmental conditions, cultural practices, and phase on which the epidemic was assessed could affect the incidence values observed. The plots studied were cultivated with a ‘Prata’ type cultivar (Pome subgroup, AAB) that has intermediate resistance to race 1 of the pathogen (Dita et al. 2018). Intermediate resistance contributes to reduce the disease progress rate and is one of the main strategies in controlling epidemics (Parlevliet 1979). The progress of FW was best described by the monomolecular model. Differences between the initial incidences were not detected and the disease rate was higher in the plot with the highest WB population. This fact indicated that the dynamics of FW epidemics may have been affected by the insect population, probably acting as dispersal agent or predisposing the banana plants to infection by *Foc* (Meldrum et al. 2013).

Aggregation was the prevalent pattern in both fields during the study. A higher degree of aggregation was observed in the plot with lower populations of WB. Higher spatial heterogeneity was observed in the managed plot compared to the plot left unmanaged. A higher degree of aggregation could be due to plant-to-plant transmission. A diseased plant acts as an inoculum source to the nearest neighbor plants. Root-to-root was the main transmission mechanism of *F. oxysporum* f. sp. *radicis-lycopersici* in tomato fields (Rekah et al. 1999). Aggregation was also the dominant t pattern detected by the geostatistical-based method, *SADIE*. The clusters of diseased plants were closer in WB-managed plots than in the unmanaged. Additionally, the degree of aggregation increased over time showing higher values in the last assessments for both methods, *D* and *I_a_*. Initial infections of stem bleeding disease of coconut palm had a random pattern in the first assessments but evolved to an aggregated pattern in the last (Carvalho et al. 2013).

At low levels of incidence, the plot managed for WB presented higher aggregation of FW. The level of aggregation was directly related to disease incidence in the plot without management. Reducing the population of a vector with management, the pathogen dispersal at greater distances is reduced and dispersal occurs likely into neighboring plants. In the umanaged plot the disease was more randomly distributed over time, with new foci appearing far from the initial foci. This may be interpreted as evidence of WB acting as a within-field vector. However, no inference can be made regarding vectoring the pathogen among neighboring banana fields.

The spatio-temporal dynamic of FW epidemics may also be impacted by the differential behavior of WB and FWB. *C. sordidu*s adults have limited mobility. Even though they have functional wings, the flight is rare, movement by crawling is limited and attracted by host volatiles (Gold et al. 2001). Unpublished studies performed in Uganda demonstrated that 81% of the *C. sordidus* released moved only up to 15 m during a six months time period. Only 3% of the individuals moved farther than 25 m (Gold et al. 2001). On the other hand, *M. hemipterus* is more active and mobile. *M. hemipterus* are good flying insects and are attracted by wounds and plant debris, but can also feed in banana plants (Fancelli et al. 2012; Molet 2013).

Although WB had limited movement, differences in spatial patterns were detected between the two fields studied and could bring new insights about the contribution of the pest in FW epidemics. *F. oxysporum* f. sp. *cubense* is a soilborne pathogen and due to the crawling habit of WB (Gold et al. 2001), soil particles can easily be found in their exoskeleton. On the other hand, FWB is a good flier and it is possible that movement in soil is more limited than WB (Molet 2013). Notwithstanding, it is expected that WB, at the individual level, had a higher potential to carry *Fusarium* propagules than FWB. It is also hypothesized that WB larvae have a preference to feed in rhizomes (Gold et al. 2001) of banana plants while larvae of FWB prefer to feed in stems or plant debris (Fancelli et al. 2012; Molet 2013). Further studies need to account for these differences in the behavior of the vector to fully understand the contributions of *M. hemipterus* in FW epidemics and test these hypotheses.

Approximately ten percent of the WB collected in fields with FW epidemics were identified carrying viable spores of *Foc* TR4 in their exoskeleton (Meldrum et al. 2013). The maximal inoculum acquisition by WB adults occurred after one hour of contact with infected pseudostems (Sánchez et al. 2021). In the environment, the contact between insects and inoculum sources, as infested soil or *Foc*-infected banana tissues, can be longer but may not impact the concentration of inoculum in their bodies. It is also of great importance the time between acquisition and inoculation of healthy plants. *Foc* propagules remained viable for approximately three days, but viability dropped exponentially after acquisition (Sánchez et al., 2021). If the infested insect does not move to new banana mats in three days (Sánchez et al. 2021), and the pathogen was not endemic in the area or widely spread in the soil, the chances of *Foc* being inoculated into healthy plants are drastically reduced.

The contribution of WB in the spatio-temporal dynamic of FW epidemics under field conditions was highlighted in this study. Fields with lower WB populations presented a more aggregated pattern, demonstrating that the dispersal of *Foc* at greater distances was minimized. These results may direct new studies to clarify the interaction between *Foc* and their potential vectors. Also, direct management practices, as rigorous WB control, to reduce the impact of FW epidemics. Better management of banana FW relies on the development of efficient strategies to reduce the dispersal of *Foc* within and between fields.

